# Reactivation of Epstein Barr Virus from Latency Involves Increased RNA Polymerase Activity at CTCF Binding Sites on The Viral Genome

**DOI:** 10.1101/2021.11.01.466781

**Authors:** Laura E. M. Dunn, Fang Lu, Chenhe Su, Paul M. Lieberman, Joel D. Baines

## Abstract

The ability of Epstein-Barr Virus (EBV) to switch between latent and lytic infection is key to its long-term persistence, yet the molecular mechanisms behind this switch remain unclear. To investigate transcriptional events during the latent to lytic switch we utilized Precision nuclear Run On followed by deep Sequencing (PRO-Seq) to map cellular RNA polymerase (Pol) activity to single-nucleotide resolution on the host and EBV genome in three different models of EBV latency and reactivation. In latently infected Mutu I Burkitt Lymphoma (BL) cells, Pol activity was enriched at the Qp promoter, the EBER region and the BHLF1/LF3 transcripts. Upon reactivation with phorbol ester and sodium butyrate, early phase Pol activity occurred bidirectionally at CTCF sites within the LMP-2A, EBER-1 and RPMS1 loci. PRO-Seq analysis of Akata cells reactivated from latency with anti-IgG and a lymphoblastoid cell-line (LCL) reactivated with small molecule C60 showed a similar pattern of early bidirectional transcription initiating around CTCF binding sites, although the specific CTCF sites and viral genes were different for each latency model. The functional importance of CTCF binding, transcription and reactivation was confirmed using an EBV mutant lacking the LMP-2A CTCF binding site. This virus was unable to reactivate and had disrupted Pol activity at multiple CTCF binding sites relative to WT virus. Overall, these data suggest that CTCF regulates the viral early transcripts during reactivation from latency. These activities likely help maintain the accessibility of the viral genome to initiate productive replication.

**Author summary:** The ability of EBV to switch between latent and lytic infection is key to its long-term persistence in memory B-cells and its ability to persist in proliferating cells is strongly linked to oncogenesis. During latency, most viral genes are epigenetically silenced, and the virus must overcome this repression to reactivate lytic replication. Reactivation occurs once the immediate early (IE) EBV lytic genes are expressed. However, the molecular mechanisms behind the switch from the latent transcriptional program to begin transcription of the IE genes remain unknown. In this study, we mapped RNA polymerase (Pol) positioning and activity during latency and reactivation. Unexpectedly, Pol activity was not enriched at the IE genes immediately after reactivation but accumulated at distinct regions characteristic of transcription initiation on the EBV genome previously shown to be associated with CTCF. We propose that CTCF binding at these regions retains Pol to maintain a stable latent chromosome conformation and a rapid response to various reactivation signals.

## Introduction

Epstein-Barr virus (EBV) is a gammaherpesvirus that establishes a persistent infection in 95% of adults worldwide. EBV maintains this persistence through a biphasic lifecycle in which it switches between latent and lytic phases. The virus establishes latency in B-cells where it exists as a circular episome and the majority of genes are believed to be epigenetically silenced via repressive chromatin and DNA methylation (1). EBV infection of B-cells leads to their transformation into immortalized lymphoblastoid cell-lines (LCLs), and is strongly associated with B-cell cancers such as Burkitt’s lymphoma and Hodgkin’s lymphoma (2).

The latent transcriptional program is highly restricted compared to the lytic program, with expression limited to a combination of six EBNA genes, two LMP genes, and multiple non-coding RNAs (3). The expression pattern of the EBNA and LMP genes determines the classification of the four distinct latency types (0-III), with 0 the most restricted (no viral protein expression) and III the least restricted (3). EBNA expression is regulated by selection from the Qp, Cp and Wp promoters (4). This promoter selection is mediated by the chromatin architecture of the EBV episome, which is dependent on host CCCTC-binding factor (CTCF) (5). CTCF is a multifunctional DNA binding protein with roles in transcriptional activation/repression, chromatin loop and boundary formation, and promoter-enhancer blocking activity (6). The EBV genome contains multiple CTCF binding sites (7), and a number of these sites have been linked to the maintenance of the latent chromatin state (8) (9) (10).

Reactivation is known to be induced by stimulation of the B cell receptor (BCR) and various stress-related signals (11). These events lead to activation of the viral Zp and Rp promoters to drive transcription of the immediate early (IE) EBV genes, BZLF1 and BRLF1, which in turn induce early lytic gene expression (12). A number of host-cell factors including BLIMP1 (13), PARP1 (10) and MYC/MAX (14) have been shown to be involved in regulating the induction of IE expression. However, the exact molecular mechanisms as to how EBV overcomes the repressive latent state to promote transcription of the lytic genes remains unknown.

EBV relies on cellular RNA polymerase II (Pol II) for transcription, although it also uses RNA Polymerase III (Pol III) to transcribe Epstein-Barr virus-encoded small RNA 1 and 2 (EBER1, 2) (15). Pol II contains an extra C-terminal domain (CTD) and the phosphorylation pattern of the CTD changes during functional switches between transcription initiation, elongation and termination (16). EBV is known to modulate Pol II transcription elongation by promoting serine 2 CTD phosphorylation to drive B-cell immortalisation. The switch from type I to III latency is regulated by Pol II pausing at the Cp promoter through association with the negative elongation factor (NELF), which facilitates recruitment of the positive transcription elongation factor (pTEFb). pTEFb maintains CTD serine 2 phosphorylation, leading to strong expression of EBV immortalization genes (17). It is not yet known if RNA polymerase pausing-elongation regulates the latent-lytic switch.

Pausing of Pol II at promoter regions, known as promoter proximal pausing (PPP), is a mechanism known to be used by herpesviruses to regulate transcription. The alphaherpesvirus, Herpes simplex virus-1 (HSV-1) utilizes PPP to regulate temporal gene expression during lytic replication (18), as does the betaherpesvirus, human cytomegalovirus (HCMV) (19). Kaposi’s sarcoma associated herpesvirus (KSHV), another gammaherpesvirus, exploits PPP to regulate the latent-lytic switch. During latency, NELF is recruited to pause Pol II on lytic KSHV gene promoters allowing for prompt expression upon stimulation of reactivation (20).

The aim of this study was to investigate changes in Pol II activity and positioning during the transition from a restricted latent state to the transcriptionally active lytic state. We hypothesized that Pol II would remain paused on the promoters of the IE genes during latency and that upon reactivation, Pol II would rapidly elongate into gene bodies. To address this hypothesis we used Precision nuclear Run On followed by deep Sequencing (PRO-Seq) (21) to map cellular Pol activity to single-nucleotide resolution on the EBV genome during latency and reactivation. As PRO-Seq does not distinguish between Pol II or Pol III we refer to all identified activity as just Pol.

In contrast to the hypothesis posed, we found no evidence of PPP on IE genes during EBV latency or reactivation. Instead, Pol activity during latency and reactivation was enriched at regions on the EBV genome that were often unassociated with known viral transcripts or promoters but correlated with CTCF binding sites. Using an EBV mutant lacking a CTCF binding site we confirmed the functional importance of CTCF binding on transcription and subsequent reactivation.

## Results

### Distribution of active polymerase on the EBV episome in latent and reactivated Mutu-I cells

To investigate the latent-lytic EBV transcriptional switch, we performed PRO-Seq on latently infected B-cells at various stages after reactivation. We first focused on the robust reactivation of Mutu-I BL cells treated with NaB/TPA. Nuclei from Mutu-I cells were harvested at 1, 4, and 12h post reactivation. The extent of RNA Pol activity during latency was first established by visualization of the data across the EBV Mutu-I episome (GenBank; KC207814.1 (22)) using the Integrative Genomics Viewer (IGV) (23). Low levels of transcriptional activity were detected across the entire EBV genome with more substantial activity at the EBERs and the BHLF1/LF3 transcripts located at the lytic origins of replication (OriLyt) (Fig. 1, top panel). We utilized the software package dREG (<u>D</u>etection of regulatory elements using <u>G</u>RO-Seq and other run-on and sequencing assays) to help identify transcription initiation regions (TIRs) based on patterns of elongating Pol identified by PRO-Seq (24). An overview of the dREG output is shown below the PRO-Seq tracks. Peaks with a false discovery rate (FDR) corrected p-value ≤ 0.05 are highlighted in green. The computed TIRs during latency were detected at the EBERs, both OriLyts, the Qp promoter and the LMP region. Except for LMP, these TIRs were consistent with the expected transcriptional pattern of type 1 latency.

**Figure 1:**
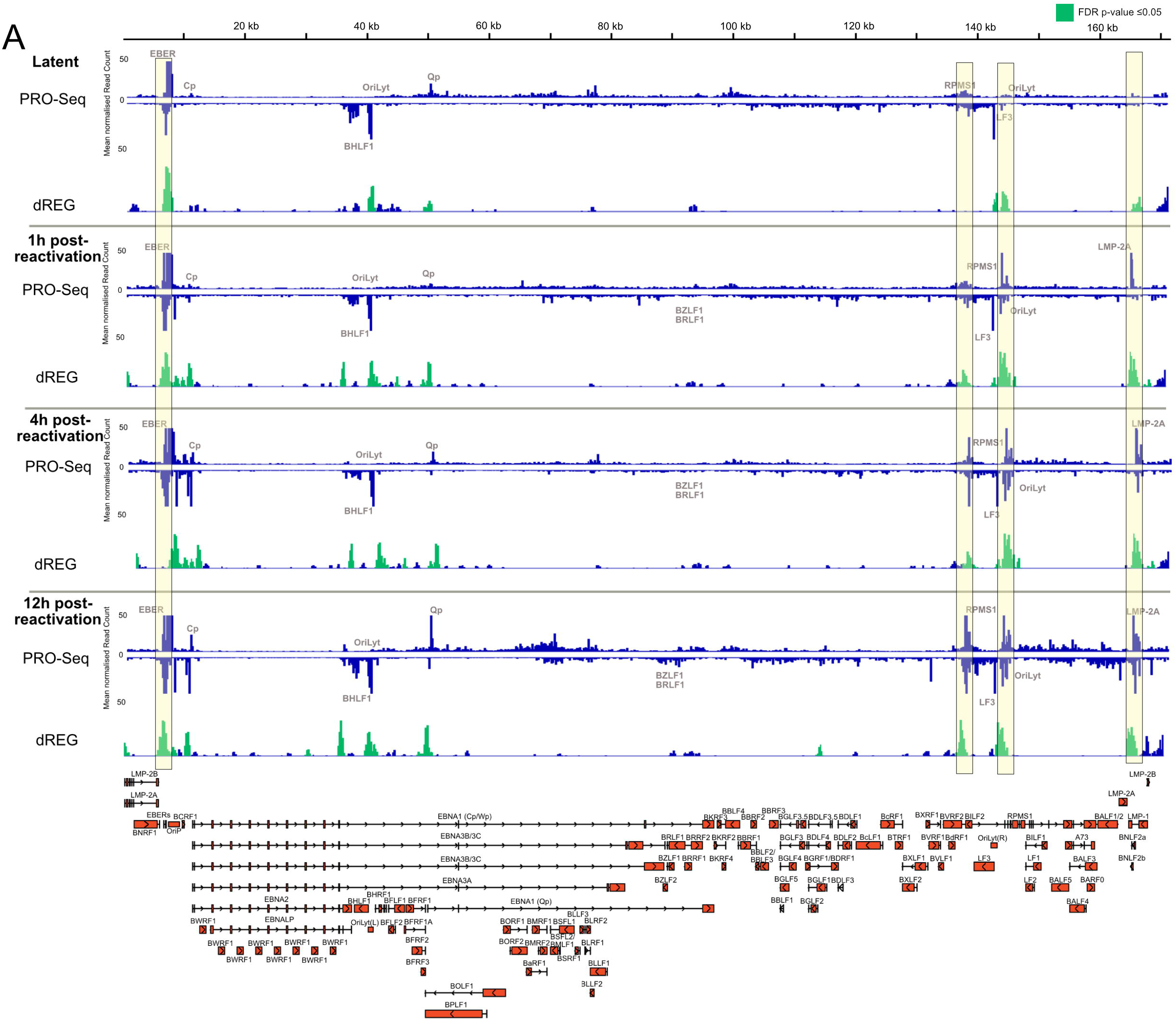
Distribution of active polymerase on the EBV episome in latent and reactivated Mutu-I cells. IGV view of the distribution of PRO-Seq reads along the EBV episome in latent Mutu-I cells and at 1, 4 and 12h post-reactivation with NaB/TPA (mean of 2-3 biological replicates, scaled relative to drosophila spike-in). dREG-called peaks from PRO-Seq, characteristic of transcription initiation regions (TIRs) (24) are shown below each PRO-Seq panel. Significant dREG peaks are highlighted in green [false discovery rate (FDR) corrected p-value ≤ 0.05]. Peaks of Pol enrichment at 1 and 4h post-reactivation are highlighted. (B-D) Volcano plots showing the DeSeq2 calculated fold changes of PRO-Seq reads for each EBV gene relative to latency at 1h post-reactivation (B), 4h post-reactivation (C) and 12h post-reactivation (D).

We next assessed the PRO-Seq transcriptional profile of EBV after reactivation. At 1 h post-reactivation, increased Pol activity relative to the latent profile developed only at the LMP-2A, RPMS1 and EBER regions, shown in highlighted regions (Fig. 1 lower panels). At 4 h post-reactivation, increased activity was limited to these same regions. By 12 h, Pol activity increased across the entire EBV genome, although predominant peaks remained at the LMP-2A, RPMS1 and EBER regions. Interestingly, most of the dREG-predicted TIRS were active during both latency and reactivation, and very few new TIRS emerged during reactivation. However, reactivation correlated with higher levels of Pol activity at these regions. The lack of Pol enrichment at the BZLF1/BRLF1 was notable, however dREG did predict a TIR at this site, but it did not pass the FDR threshold. Expression of these genes by 24 h post-reactivation was confirmed by RT-qPCR (Fig. S1A), indicating a lack of activity at this region was not due to inefficient reactivation.

### Distinct patterns of transcriptional enrichment during EBV reactivation are associated with CTCF binding sites in Mutu-I cells

In experimental support of the dREG computed TIRs, we used data from a previous Assay for Transposase-accessible Chromatin Sequencing (ATAC-Seq) experiment (GEO: GSE172476) in 2 sub-populations of latent Mutu-I cells, MUN14 and M14, and aligned this to the dREG data (Fig. 2A). TIRs at the EBERs, BHLF1, Qp, OriLyt(R) and LMP-2A showed an alignment to ATAC-Seq peaks, indicating that these predicted TIRs are located at sites of accessible chromatin. As TIRs discovered by the dREG package have been shown to be associated with sites of transcription factor binding (24), we next asked whether the TIRs identified on the EBV episome were linked to transcription factor binding sites. We utilized the EBV portal database (7) to align transcription factor ChIP-Seq data to the Mutu-I latent and reactivated dREG-called peaks generated from the PRO-Seq data. The most striking observation was that for virtually every dREG site, a CTCF peak also aligned (Fig. 2A).Though not every CTCF site aligned to a TIR, the strong correlation warranted further investigation. We therefore examined some of the TIRs at higher resolution using IGV. Commonly, peaks of Pol activity were visible next to or over sites of CTCF binding, and often on both strands of the genome. This is shown in detail at the EBER region in Fig. 2B, with extensive Pol activity on both strands over and adjacent to the CTCF site during reactivation. The CTCF site upstream of the Cp promoter is visible in the same figure, with bidirectional Pol activity over the CTCF site upon reactivation. The BHLF1/OriLyt(L)/Qp region is shown in Fig. 2C, with 3 CTCF sites coinciding with enriched Pol activity. Further examples of extensive Pol activity adjacent to and flanking CTCF sites during reactivation are shown for the RPMS1/OriLyt(R) region (Fig. 2D) and the LMP-2A region (Fig. 2E). The CTCF binding site at BZLF1 is shown in Fig. 2F. An enrichment of Pol activity can be seen over this site at 12 h post-reactivation, suggesting it is more relevant later in reactivation compared to other CTCF binding sites.

**Figure 2:**
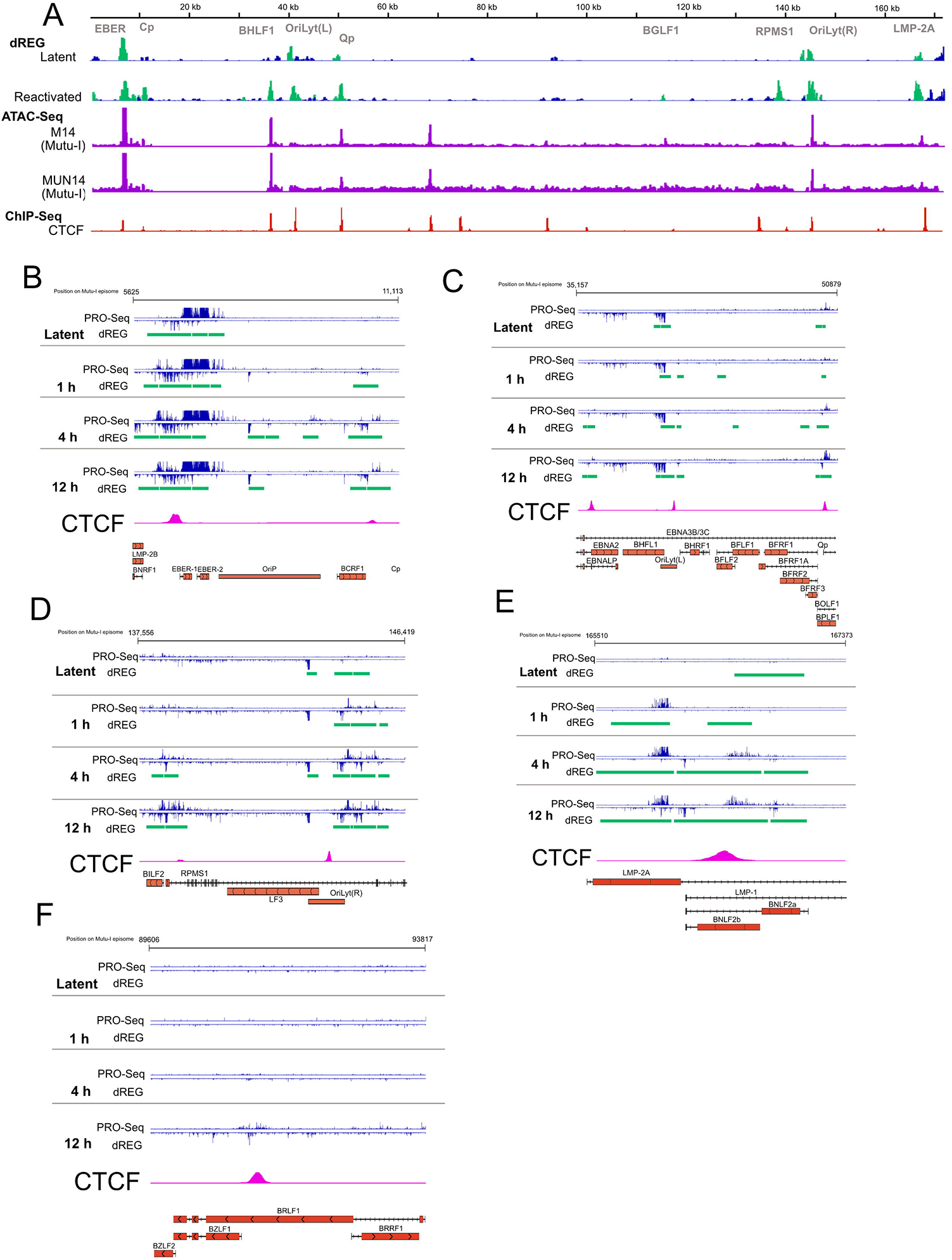
PRO-Seq identifies distinct patterns of Pol activity on the EBV episome during reactivation associated with CTCF binding sites. (A) Alignment of dREG-called peaks in latent and reactivated Mutu-I cells, ATAC-Seq data from 2 sub populations of latent Mutu-I cells and CTCF ChIP-Seq. (B-F) High resolution IGV views of PRO-Seq tracks from EBV genome aligned with dREG-called TIRs and regions of CTCF binding at EBER – Cp region (B), OriLyt (L) – Qp region (C), RPMS1 – OriLyt(R) region (D). LMP-2A exon 1 region (E), or BZLF1-BRLF1 region (F). The data shown are means of 2-3 biological replicates, scaled relative to drosophila spike-in. In genome annotation, black bars indicate known TSSs or TTSs, with coding regions shown in red and the orientation of the gene is indicated by arrows.

Viewing the data at this high resolution also indicated that the PRO-Seq peaks and predicted TIRs did not map near known EBV gene transcription start sites (TSSs) or promoters suggesting that promoter proximal pausing of Pol was not a major feature of latency or reactivation. For example, the increased activity associated with the EBERs was found upstream of EBER-1, on both strands of the genome in an area encoding no known transcripts (Fig. 2B). A further example was noted at LMP-2A, where the increase in Pol occurred over the exon downstream of the start codon, with additional activity appearing later in reactivation 150 bp (on the opposite strand) and 500 bp downstream of the initial peak (Fig. 2E). The lytic EBV transcriptome is known to be more complex than the GenBank annotation (26,27). However, even when novel lytic transcripts (using transcript information from O’Grady et al. (26)) were included in the alignment, the majority of PRO-Seq peaks did not align to canonical or novel TSSs (Fig. S2A-F). Taken together, the PRO-Seq data has indicated that enrichment of Pol activity during EBV reactivation is inconsistent with promoter proximal pausing upstream of known EBV genes and is instead associated with CTCF binding sites.

### Pol enrichment at CTCF binding sites is a consistent feature of EBV reactivation in multiple cell-types

To assess whether these TIRs were specific to Pol activity in Mutu-I cells and the NaB/TPA method of reactivation, we repeated the experiment using another EBV positive Burkitt’s lymphoma cell-line, Akata, reactivated with anti-IgG. It should be noted that the number of EBV genomes in Akata cells is lower than in Mutu-I cells, causing the Akata PRO-Seq data to comprise approximately 10-fold lower normalized EBV aligned reads (Table S1, S2). As a result, it was difficult to visualize distinct changes in PRO-Seq activity on IGV until 12 h post-reactivation, when transcription was more robust (Fig. S3A). We confirmed this was not due to inefficient reactivation as BZLF1 was detectable by RT-qPCR at just 4 h post-reactivation (Fig. S1B). However, using dREG we were able to identify relatively enriched regions of Pol activity, which again showed a strong association to CTCF binding sites throughout latency and reactivation (Fig. 3A).

**Figure 3:**
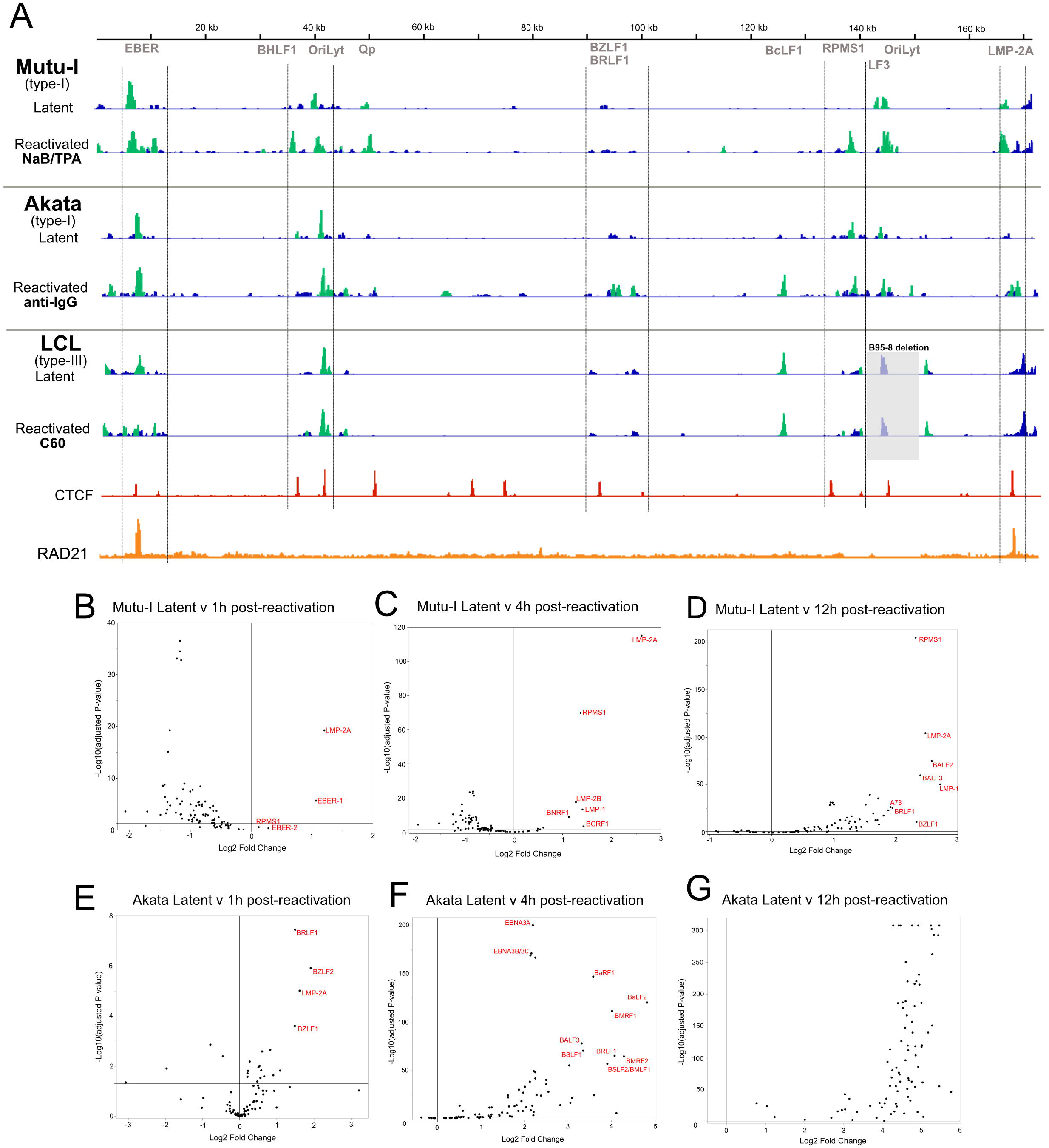
Sites of Pol enrichment during latency and reactivation in multiple models of EBV reactivation align with CTCF binding sites. (A) Summary of dREG-called peaks in latent and reactivated Mutu-I cells, Akata cells and LCLs, aligned to CTCF and RAD 21 binding sites. Hatched areas indicated regions that were consistent in all cell lines. Significant dREG peaks are highlighted in green (false discovery rate (FDR) corrected p-value ≤ 0.05). For Akata and Mutu-I cells, all reactivated timepoint tracks are overlaid. (B-F) Volcano plots showing the DeSeq2 calculated fold changes of PRO-Seq reads for each EBV gene relative to latency at Mutu-I 1h post-reactivation (B), Mutu-1 4h post-reactivation (C), Mutu-I 12h post-reactivation (D), Akata 1h post-reactivation (E), Akata 4h post-reactivation (F), Akata 12h post-reactivation (G),

Despite both systems sharing type 1 latency, some variations in the locations of the significant dREG-called TIRs were observed between Mutu-I and Akata cells. However, most of these cell type-specific peaks also aligned near CTCF binding sites. Similar to Mutu-I, positions of most dREG-predicted TIRs in Akata were unchanged from latency, even by 12 h when transcriptional activity was extensive. Interestingly, a peak at the promoter of the transcriptional activator, BRLF1, appeared in Akata during reactivation (Fig. 3A, middle panel). Transcription at this promoter generates a bicistronic mRNA containing BZLF1 and BRLF1 and these genes are separated by a CTCF site. The early appearance of a peak in the region containing the major lytic transcriptional activators in Akata and not Mutu-I suggests that these cells may differ in their kinetics of reactivation. This was supported by the DeSeq fold change quantification of EBV gene reads, with more viral genes significantly increasing in Pol activity at 1h and 4h post-reactivation, including BRLF1 and BZLF1, in Akata cells relative to Mutu-I (Fig. 3B-G). Supporting what was visible on the PRO-Seq tracks (Fig. 1), significant increases on Mutu-I at 1 and 4h post-reactivation were mostly limited to genes in the RPMS1-LMP region (Fig. 3B, C) and BZLF1 and BRLF1 were only significantly increased in Pol activity at 12h post-reactivation (Fig. 3D).

We next used PRO-Seq to investigate reactivation in another reactivation model: type-III latency LCLs reactivated with the small molecule C60 (28). Cells were treated with either DMSO (in which the genome remains in a latent state) or C60 treatment to induce reactivation for 24h and then PRO-Seq. After 24h of treatment with C60 to induce reactivation, lytic gene activity increased relative to cellular control gene GUSB (Fig. 5B-F). No viral genes had statistically more active Pol, relative to DMSO treatment (Appendix 2). However, consistent with the other cell types studied, the dREG-predicted TIRs aligned with CTCF binding sites (Fig. 3A, lower panel), except for a peak at BcLF1 -which was also found in Akata cells. As with both Mutu-I and Akata cells, clusters of TIRs were identified at the EBERs, Orilyt(L) and RPMS1 throughout latency and reactivation. A peak was called at the LMP region but did not reach the FDR significance threshold. An overview of dREG-called TIR peaks from all three cell-types aligned to CTCF binding sites is shown in Fig. 3, with hatched areas highlighting regions that were consistent in all models. CTCF commonly functions in complex with cohesin. Therefore, a track containing ChIP for the cohesin subunit RAD21 was also included, and peak binding aligned with dREG peaks at the EBER and LMP regions.

### CTCF/cohesin rearrangements occur during EBV reactivation

Using three different models of EBV reactivation we have shown that Pol activity during reactivation was enriched at CTCF binding sites. However, it remained unknown whether CTCF is directly involved or if increased transcriptional activity was an indirect result of genome accessibility. To investigate the role of CTCF binding on the EBV episome during early reactivation, ChIP-qPCR was performed on latent (untreated) and 4 h reactivated Mutu-I and Akata cells. Pol II ChIP confirmed an increase in Pol II on the EBV genome during reactivation (Fig. 4A, D). At 4h post-reactivation in Mutu-I cells, there was a decrease in CTCF binding at all 3 sites examined on the EBV genome (LMP-CTCF, Cp-CTCF, Qp-CTCF) (Fig. 4B), suggesting CTCF binding decreases upon reactivation. However, ChIP-qPCR in Akata cells gave contrasting results and indicated increased CTCF at the LMP-CTCF and Qp-CTCF binding sites but was unchanged at the Cp-CTCF site (Fig. 4E).

**Figure 4:**
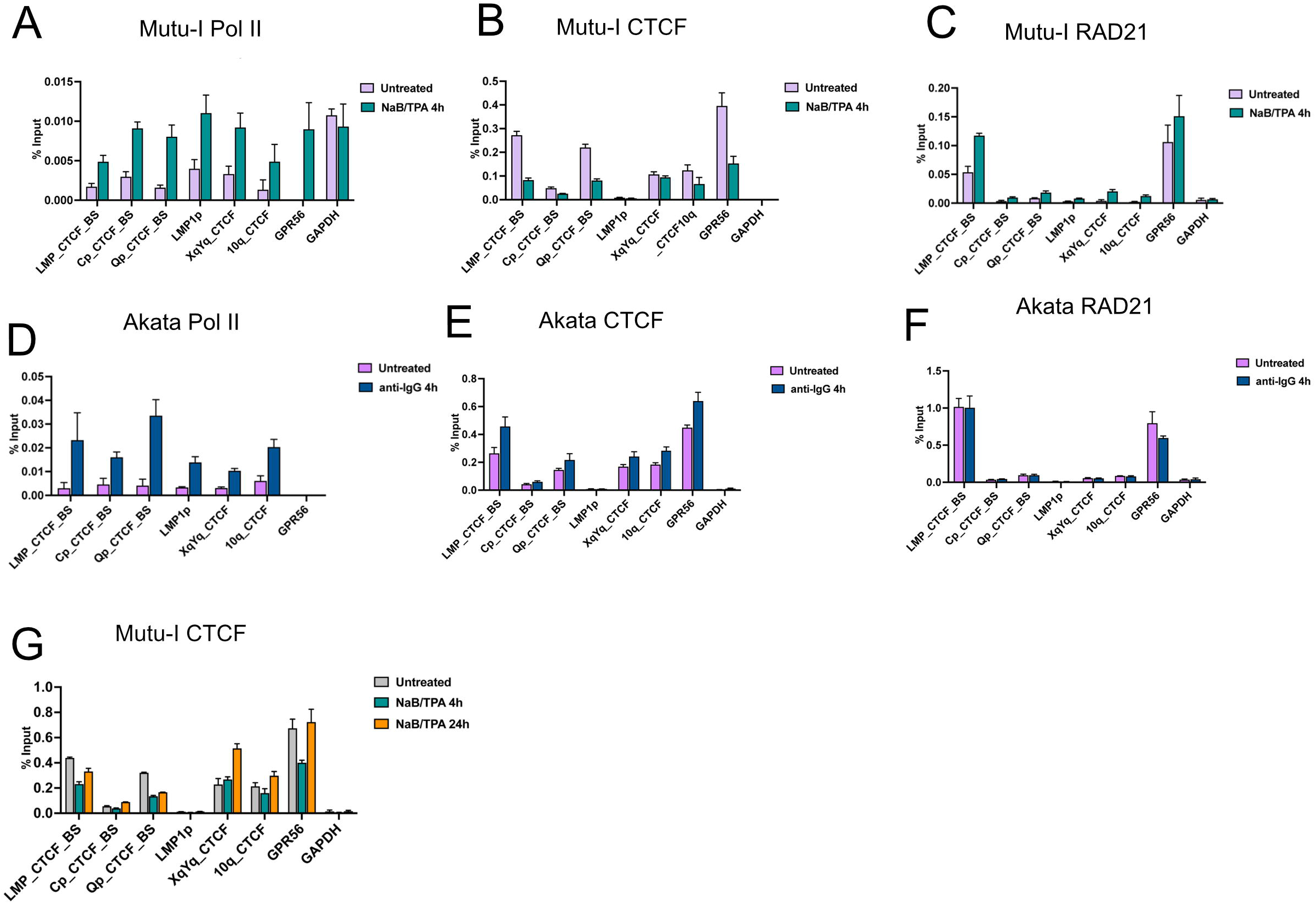
CTCF/cohesin binding rearrangements during EBV reactivation. ChIP-qPCR at CTCF binding sites on the EBV and human genomes in latent and reactivated Mutu-I cells using antibodies indicated. (A) Mutu-I Pol II (B) Mutu-I CTCF (4h) (C) Mutu-I RAD21 (D) Akata Pol II (E) Akata CTCF (F) Akata RAD21 and (G) Mutu-I CTCF (24h)

**Figure 5:**
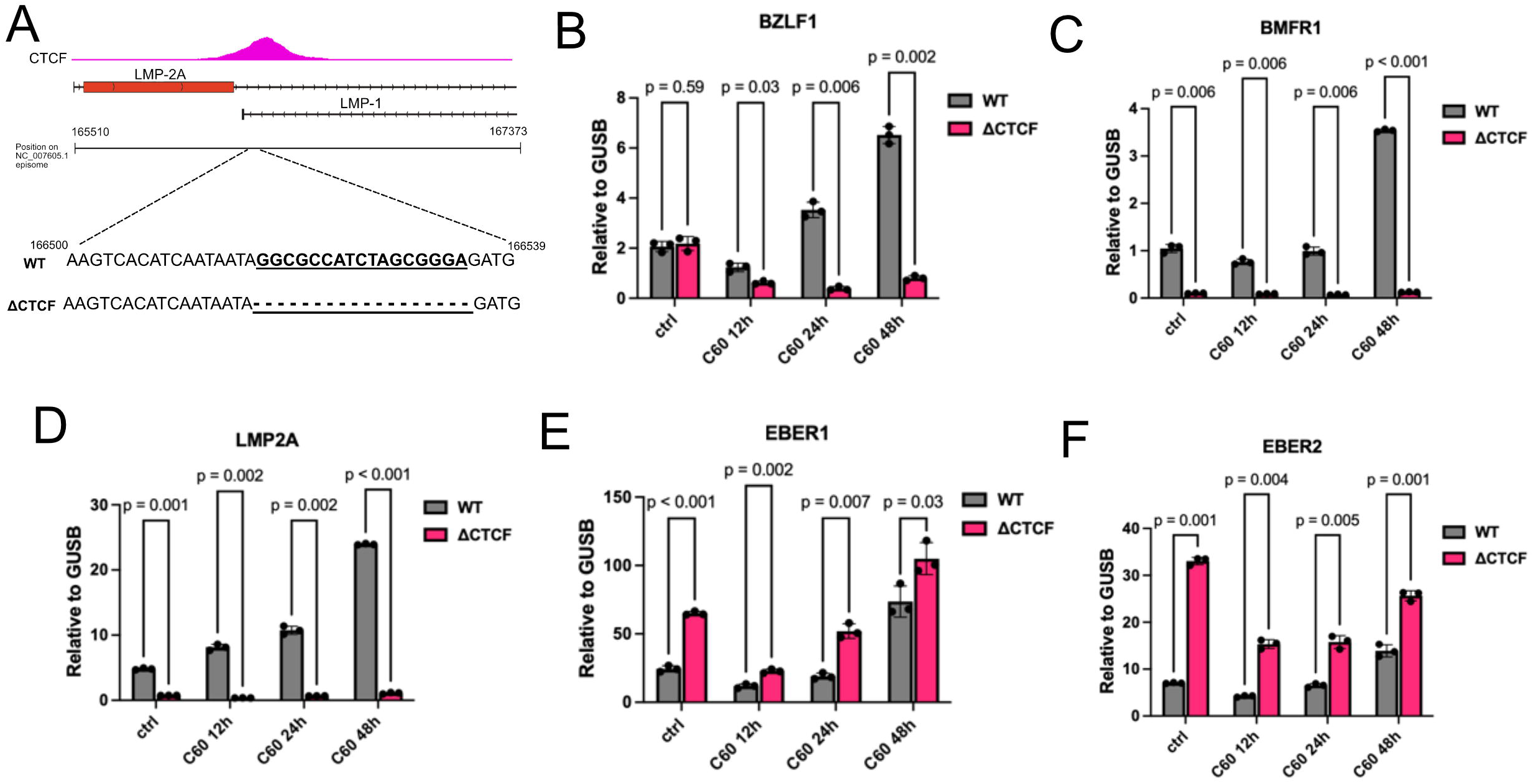
An EBV mutant with a mutated CTCF binding at LMP does not reactivate from latency. (A) Schematic diagram of the deletion in the ΔCTCF EBV mutant. B-F. RT-qPCR for EBV genes BZLF1(B), BMRF1(C), LMP-2A(D), EBER-1 (E) and EBER-2 (F) after induction of reactivation with the small molecule, C60 at 12, 24 and 48 h post-treatment of LCL WT or ΔCTCF. Statistical significance was determined by students t-test.

ChIP-qPCR for the cohesin subunit RAD21 in Mutu-I cells revealed an increase in binding at 4h post-reactivation at the LMP region (Fig. 4C) but was unchanged in Akata cells (Fig. 4F). This suggested that CTCF/cohesin complex is dynamic and rearranged upon reactivation, but the details of this process are likely dependent on cell-type and reactivation stimulus. Our PRO-Seq data indicated Mutu-I cells have slower reactivation kinetics than Akata cells, and repeating ChIP-qPCR for CTCF in Mutu-I cells at 24h post-reactivation revealed CTCF binding increased relative to 4h (Fig. 4G). This further supports the conclusion that CTCF rearrangements are linked to the kinetics of reactivation.

### A mutation that disrupts CTCF binding at LMP alters transcriptional activity and precludes reactivation

To further investigate the functional role of CTCF during EBV reactivation we utilized the LCL EBV mutant (ΔCTCF) which contains an 18bp deletion at the LMP-2A CTCF-binding site (Fig. 5A) resulting in a disruption of CTCF binding (9). The ability of this mutant to reactivate was first assessed using RT-qPCR for viral genes after treatment with C60. The ΔCTCF mutant was found to have a limited ability to reactivate from latency, with very low expression of the lytic genes BZLF1 and BMRF1, even by 48 h post-reactivation when the WT exhibited robust expression (Fig. 5B, C). In contrast to WT, ΔCTCF was also unable to express LMP-2A throughout latency and reactivation (Fig. 5D) and showed dysregulated expression of both EBER-1 and EBER-2 (Fig. 5E, F). These results implicate the importance of CTCF binding in the regulation of EBV transcription because disruption of only a single CTCF site precluded expression of lytic mRNAs and reactivation. Therefore, we used PRO-Seq to investigate transcriptional activity in detail on the ΔCTCF mutant 24 h after treatment with either DMSO or C60, compared to the previously studied WT LCL. Compared to the PRO-seq profiles of Mutu-I and Akata cells shown previously, higher activity across the LCL genome (Fig. S3B) was expected because these cells maintain type III latency – the most transcriptionally active of all EBV latency types (3).

Initially focusing on latency (DMSO treatment), the most striking difference between ΔCTCF and WT was a decrease in transcriptional activity (approx. 2.7x (Table S2)) across the mutant genome. To better illustrate differences between the two transcriptional profiles, a heatmap of the PRO-seq data was generated to visualize the density of reads on both DNA strands (Fig. 6A). This heatmap confirmed the initial PRO-Seq analysis, and indicated that ΔCTCF had a lower level of Pol activity across most of the genome relative to WT, although specific sites of the mutant genome had higher activity than WT. Most noticeable of these was an increase in activity at ∼11 kb on the ΔCTCF genome (near the Cp promoter), continuing ∼30 kb downstream through the W repeat region. Visualization of the PRO-Seq data at this region using IGV (with adjusted scaling to equate total viral reads between viruses) revealed strong peaks on the ΔCTCF genome at the Cp promoter and over the CTCF site just upstream of Cp (Fig. 6B). The EBERs also increased in activity in ΔCTCF relative to WT and are flanked by a CTCF site (Fig. 6B). There were differences in activity at the Qp promoter (50kb), with ΔCTCF exhibiting a reduction in several prominent peaks visible in WT (Fig. 6C). Further differences were seen at ∼139 kb (RPMS1); in particular, Pol activity increased over and adjacent to the CTCF site in the mutant, primarily on the minus strand where no genes map (Fig. 6D). A clear shift in Pol enrichment between the viruses was seen at the LMP region. The WT displayed strong peaks of bidirectional transcription over the CTCF site at LMP-2A (site of the deletion in the mutant) (Fig. 6E). ΔCTCF had a reduction in activity at this site with a marked increase downstream over LMP-2B.

**Figure 6:**
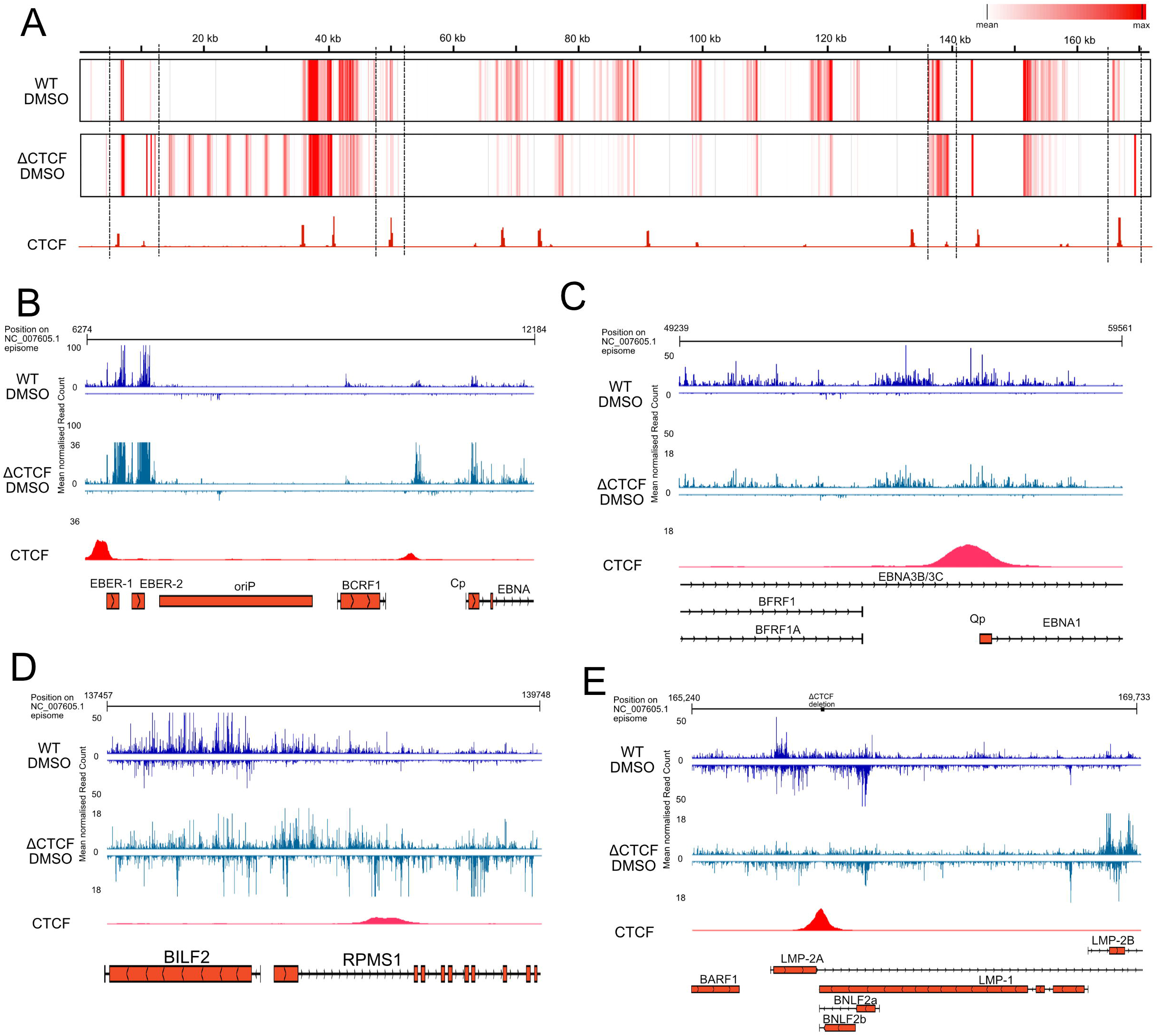
Mutation of CTCF binding site at LMP alters transcriptional activity at other CTCF binding sites during latency. (A) Heatmap of Pol density on both strands of the viral genome, in 50 bp bins. The background level was determined for each virus as the mean of each 50 bp window and regions in red indicate sites of Pol enrichment above this level. (B-E) High resolution IGV view of PRO-Seq tracks from EBV genome regions with disrupted activity in the ΔCTCF mutant during latency, aligned to CTCF binding sites at the EBER – Cp region (B), Qp region (C), RPMS1 region (D), or LMP region (E).

After 24h of treatment with C60 to induce reactivation, specific sites of transcriptional enrichment were identified by fold change analysis across the viral genome and indicated differences between WT and ΔCTCF at the Cp, LMP, BHRF1 and BZLF1 regions (Fig. S3B). Closer analysis of the Cp region of WT revealed 3 peaks of increased Pol activity after C60 treatment at BCRF1, over the CTCF binding site, and at the Cp promoter (Fig. 7A). The peak activity aligning to BCRF1 and Cp promoter were located downstream of the start codon rather than directly over promoter regions. Spline interpolation analysis indicated that the peak increases directly over the CTCF binding site Cp promoter were statistically significant (in highlighted regions) after reactivation in WT (Fig. 7A). In ΔCTCF, although there was a significant increase at the Cp promoter, there was no change the CTCF binding site and a complete lack of activity at BCRF1. These differences in Pol activity at the Cp region correlated with altered Pol activity throughout the W repeats, which was heavily restricted in WT but highly active in ΔCTCF (Fig. S3B).

**Figure 7:**
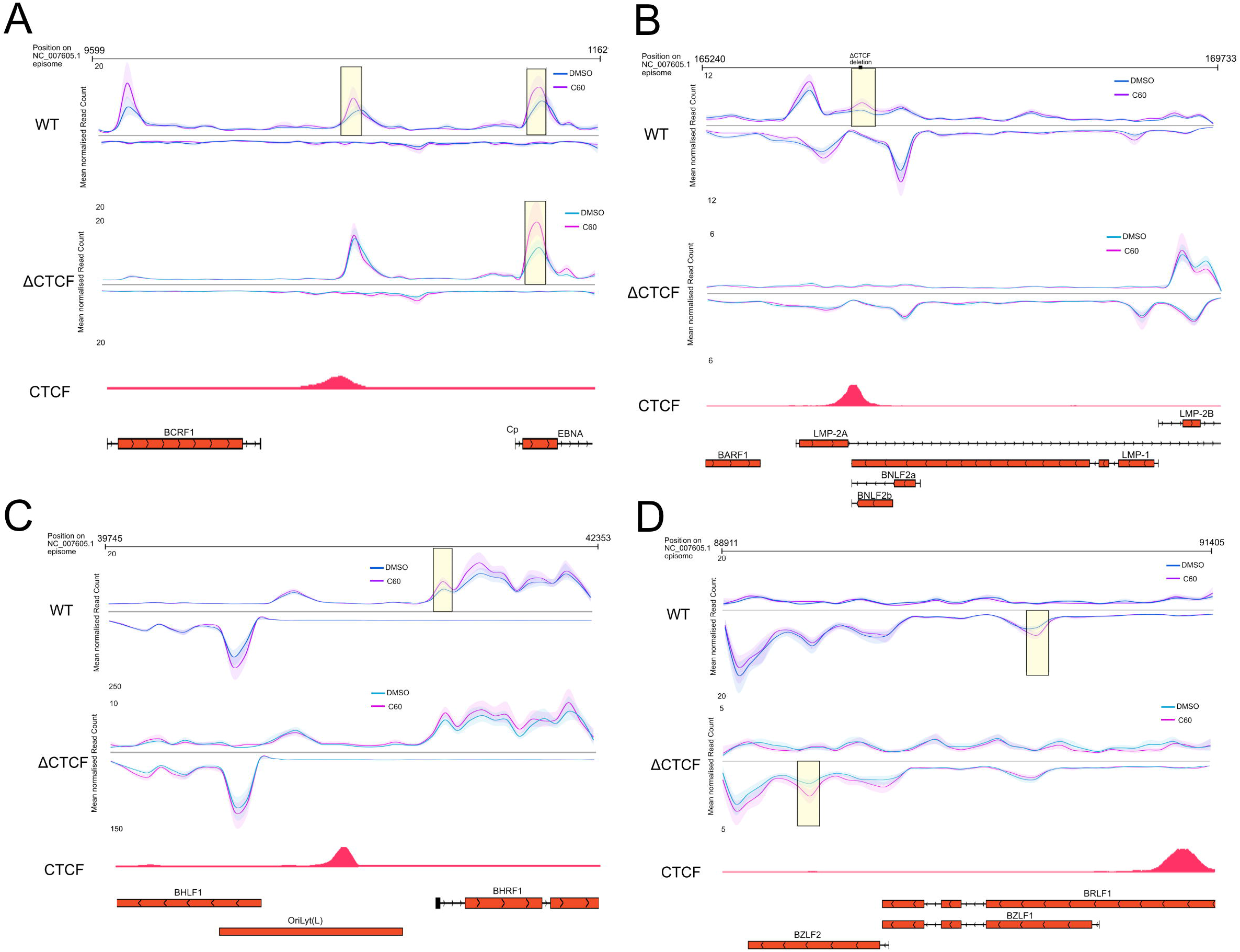
A mutation of the CTCF binding site at LMP changes transcriptional activity at other CTCF binding sites during stimulation of reactivation. (A-D) Spline interpolation of PRO-Seq reads from EBV genome regions with disrupted activity in the ΔCTCF mutant during reactivation, aligned to CTCF binding sites at BCRF1 – Cp region (A), LMP-2A region (B), BHLF1-BHRF1 region (C) and BRLF1-BZLF1 region (D). The shaded area shows confidence of fit. Highlighted regions indicate regions of statistical significance between DMSO and C60 treatment.

At LMP-2A, transcriptional activity was significantly increased over the CTCF binding site after reactivation in WT (Fig. 7B). However, in ΔCTCF, there was no activity at this site in either DMSO and C60 treatment and as previously noted, a subsequent peak at LMP-2B, but this was unchanged after C60 treatment. At BHRF1, approximately 500bp downstream of the CTCF site, and which flanks the highly active BHLF1 transcript at OriLyt(L), C60 treatment led to significant increases of Pol activity at the BHRF1 promoter region in WT (Fig. 7C). Such differences were less striking in ΔCTCF and did not reach statistical significance. Activity at the BZLF1 region was also altered between viruses with a significant increase in Pol activity in WT after C60 treatment in the gene body of BZLF1, ∼700bp downstream of the CTCF binding site (Fig. 7D). In ΔCTCF, this increase in Pol activity appeared to shift a further ∼900bp downstream, over the gene body of BZLF2.

In summary, these analyses have indicated that peaks of transcriptional activity occur adjacent to, or over multiple CTCF sites during LCL latency. Aberration of a single CTCF binding site at the LMP region disrupts transcriptional activity at other CTCF sites during LCL latency and this phenotype correlates with an inability to reactivate from latency. These data support the previous findings in Mutu-I and Akata cells and provide functional evidence of the importance of CTCF binding in the control of EBV transcription during latency and reactivation.

CTCF is known to regulate 3D genome interactions, and Hi-C analyses for both latent LCLs and Mutu-I cells have recently been reported (42). We therefore aligned Hi-C DNA interactomes of LCLs and Mutu-I with our ProSeq data (Fig. 8). This overlay reveals that the sites of transcriptional enrichment identified here by PRO-Seq in both Mutu-I and LCLs aligned with intragenomic contact sites identified in the Hi-C study (Fig. 8) (42). The interaction site deleted in ΔCTCF (LMP1/2 region) is highlighted (9), and clearly shows that this region makes multiple contacts with other regions of the viral genome, especially the EBER-oriP region. In LCLs, additional contacts are detected with Qp, BZLF1and RPMS1, suggesting the CTCF interactome on the viral genome provides coordinate control of viral reactivation.

**Figure 8:**
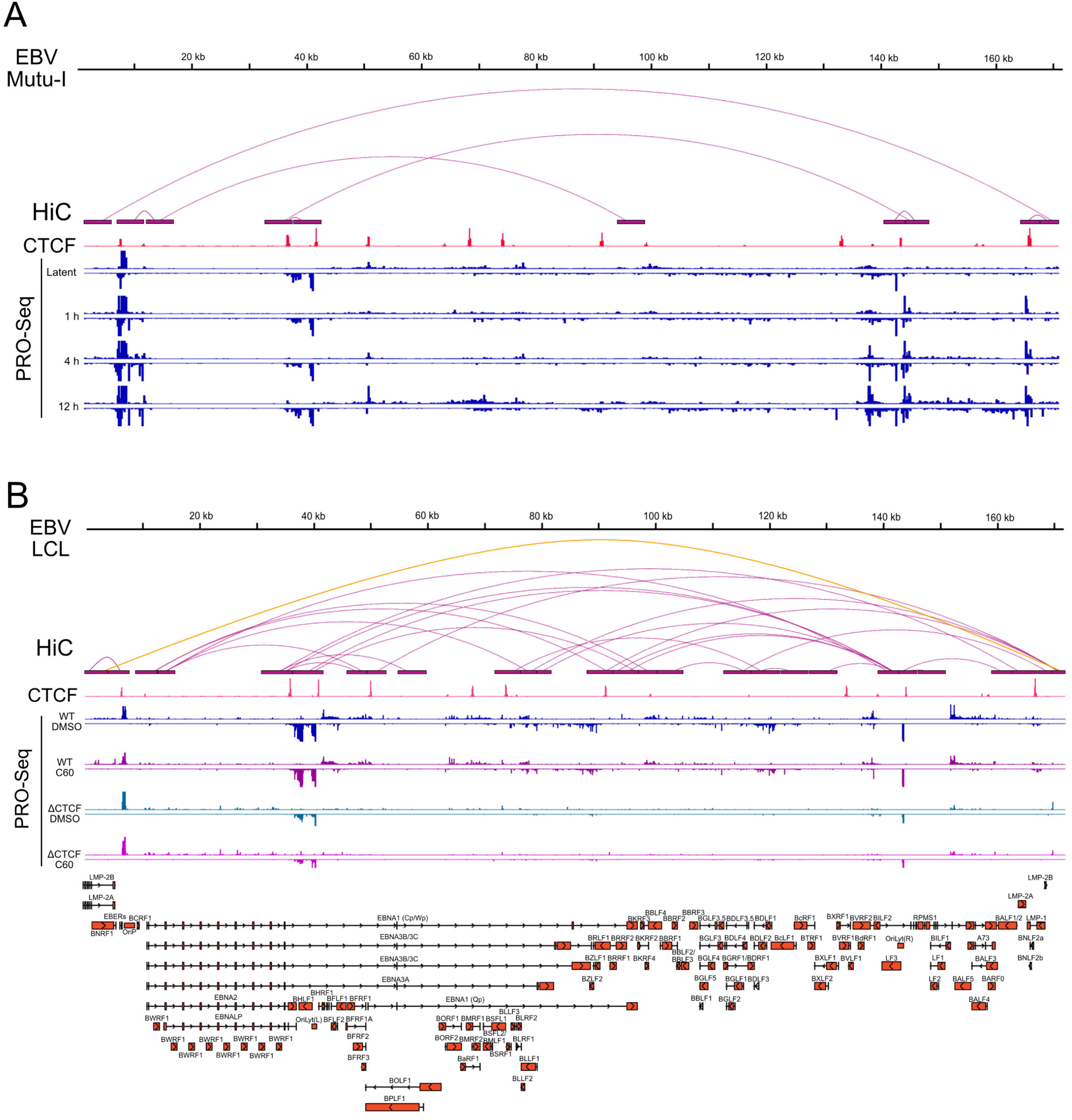
Alignment of PRO-Seq and previously obtained HiC data. Significant chromatin loops identified from HiC in Morgan et al., (42) were aligned to PRO-Seq data from (A) latent and reactivated Mutu-I cells and (B) latent and reactivated WT and ΔCTCF LCLs. The orange loop highlights an interaction found to be lost in ΔCTCF in Chen et al., (9).

## Discussion

We initially speculated that EBV regulates the latent to lytic switch at the elongation step of transcription via release of Pol II from PPP. However, PRO-Seq data from EBV Mutu-I latency (Fig. 1A) showed no evidence of PPP on any EBV genes normally expressed during productive infection. Instead, the data revealed a background level of transcriptional activity across the entire viral genome, indicating that transcription during latency is not entirely silent – a notion that is also becoming more apparent in studies of other herpesviruses such as HSV-1 (29). We should note that PRO-Seq only measures transcriptional activity, and the ultimate fate of a transcribed RNA depends on its stability, processing, interactions, polyadenylation and export to the cytoplasm. Nevertheless, peak regions of activity revealed in the PRO-Seq profiles were consistent with what was expected in type I latency (30)(31).

After Mutu-I reactivation, regions of increased Pol activity did emerge but were rarely located at promoters of viral genes, even after alignment to novel transcript data. These transcriptional increases were predominantly located on both strands of the genome around the EBERs, RPMS1 and LMP2A – all traditionally latency-associated. It was notable that the IE transcriptional activators, BZLF1 and BRLF1, did not exhibit significant increases in Pol activity until 12h post-reactivation in Mutu-I cells. EBV reactivation may potentially be analogous to biphasic HSV-1 reactivation, which begins with widespread transcriptional de-repression across the viral genome, allowing for accumulation of the IE transcriptional activator VP16 (32). It is therefore possible that the high levels of activity during EBV reactivation are sites of transcription initiation, as indicated by computation using dREG, and that final gene expression is regulated through other co-or post-transcriptional mechanisms, leading to BZLF1 accumulation. This is supported by previous findings indicating that the EBV transcriptome is highly complex, with abundant antisense transcription, multiple splice junctions, alternative promoter usage and readthrough transcription (26,33).

dREG predicted TIRs can also indicate sites of enhancer RNA (eRNA) initiation and therefore lytic gene transcription could be stimulated through long-range interactions between these potential enhancer regions and the IE gene promoters (34). We also cannot rule out that the peaks represent Pol III activity because PRO-Seq clearly detected the strong Pol III EBER transcription and dREG has been shown to detect Pol III TIRs (24). This may also explain the apparent lack of PPP, as Pol III has a greatly reduced pause time relative to Pol II (35)

We discovered a strong correlation between Pol activity and CTCF binding sites on the EBV genome, during both latency and reactivation. It was notable that all cell-lines used in this study had well defined PRO-Seq peaks during latency at the EBERs and BHLF1– both of which are located adjacent to CTCF binding sites. Upon reactivation, sites with enriched transcriptional activity, characteristic of initiation, were consistently located adjacent to or over CTCF binding sites in all experiments. CTCF has previously been linked to transcription initiation on the human genome (36), and promoter-proximal CTCF-CTCF loops have been shown to be directly associated with promoter-enhancer contacts to regulate transcription (37). Recent studies have also indicated that active transcription is required to maintain these enhancer-promoter contacts (38).

CTCF is known to be important for maintenance of the epigenetic state during EBV latency (8–10) and so it is perhaps unsurprising that it is also involved in reactivation. Our ChIP data indicated CTCF binding is altered during reactivation, however, this contrasts with a previous study that stated CTCF binding is unchanged during EBV reactivation (39). A possible explanation for this difference is that we looked at an earlier time point, 4h, compared to 24h in the previous study. Indeed, ChIP-qPCR at 24h post-reactivation in Mutu-I cells showed a recovery in CTCF binding at both the LMP and Cp CTCF loci (Fig. 4G). It is also likely linked to the cell/virus background and method of reactivation as the previous study used Akata cells, which we found to give contrasting results to Mutu-I. In addition, the previous study primarily focused on the BZLF1 CTCF site which we found to align less well with altered transcriptional activity during reactivation compared to other CTCF sites.

CTCF is involved in maintaining the distinct chromatin loops found in different EBV latency stages (5) and therefore its role in reactivation is likely linked to alteration of viral genome conformation. CTCF function in loop formation involves cohesin, which forms a ring structure around the DNA (40). ChIP for the cohesin subunit, RAD21, revealed peaks at the EBER and LMP regions (Fig. 3A), and these regions are known to physically interact in DNA loops (7). Interestingly, Pol activity at both regions was increased during reactivation in all 3 of the cell-lines used in the current study. ChIP-qPCR for RAD21 in Mutu-I cells revealed an increase in binding at 4h post-reactivation at the LMP region, contrasting with the CTCF result. This is possibly due to the unstable nature of the CTCF/cohesin complex (41). Overall, these data suggest that CTCF/cohesin binding is dynamic and rearranges upon reactivation, but the details of this process are linked to cell-type, the reactivation method, and the subsequent kinetics of reactivation.

The functional link between CTCF binding and reactivation was confirmed using a ΔCTCF (at LMP locus) EBV mutant. Aside from an overall lower level of Pol activity, the differences in transcriptional activity between WT and ΔCTCF were mostly confined to alterations at or near CTCF binding sites during latency. These differences correlated with ΔCTCF being unable to reactivate and express lytic viral mRNAs. The ΔCTCF virus has already been shown to lose an interaction upstream of the OriP region, which is flanked by two CTCF sites at the EBERs and Cp (9). Interestingly, PRO-Seq of the ΔCTCF virus showed that transcription was dysregulated at these regions relative to WT, suggesting that the transcriptional differences resulted from CTCF-mediated variation in genome conformation. Another ΔCTCF EBV mutant, with a deleted binding site at Qp, also displayed altered transcriptional patterns (8). Hi-C analysis of both latent LCLs and Mutu-I cells has indicated that disrupting PARP1-mediated stabilization of CTCF on the viral genome leads to alternative genome conformations (42). In addition, viral gene expression was shown to be dysregulated in these LCLs and the regions with altered genome conformations strongly correlated with the dysregulated viral genes. Notably, the sites of transcriptional enrichment identified here by PRO-Seq in both Mutu-I and LCLs aligned with intragenomic contact sites identified in the Hi-C study (Fig. 8) (42). Although some sites with transcriptional changes do not loop with LMP, it is apparent that the 3D structure of the EBV genome is complex, increasing the likelihood that disruption of certain interactions may disrupt others indirectly. Taken together, these studies support a link between transcription, CTCF binding and the structure of the EBV genome.

In summary, our data suggests that upon reactivation signals, CTCF/cohesin binding is rearranged to alter the 3D structure of the viral genome. We hypothesize that this increases accessibility of Pol and other factors to the viral genome to increase Pol activity proximal to CTCF sites. High levels of activity at these regions may aid maintenance of the open, accessible chromatin state during the productive phase of infection.

## Materials and methods

### Cells

EBV positive Burkitt’s lymphoma cell line Mutu I and Akata were maintained in RPMI 1640 containing 12% FBS and antibiotics (penicillin and streptomycin). Mutu I was obtained from Dr. Jeffrey Sample, Penn State University Hershey Medical School, PA. Akata was obtained from Dr. Elena Mattia, University of Rome SAPIENZA, Italy. The EBV WT and ΔCTCF LCLs have been previously described in (9). *Drosophila melanogaster* S2 (ATCC) cells were grown in Schneider’s medium (Lonza) containing 10% foetal bovine serum (FBS) and maintained at 23°C

### EBV reactivation

For PRO-seq, Mutu I cells were treated with NaB (1 mM) and TPA (20 ng/ml) for 0, 1, 4, or 12 hrs, Akata cells were treated with 10 µg/ml of goat anti-human IgG (Sigma I1886) for 0, 1, 4, or 12 hrs and the EBV WT and ΔCTCF LCLs were treated with DMSO or C60 (5 μM) for 24h. For ChIP assays, Mutu I cells were untreated or treated with NaB (1 mM) and TPA (20 ng/ml) for 4 or 24 hrs, and Akata cells were untreated or treated with 10 µg/ml of goat anti-human IgG for 4 hrs. For RT-qPCR, the EBV WT and ΔCTCF LCLs were treated with DMSO or C60 (5 μM) for 12, 24, and 48h.

### Nuclei isolation

Nuclei were isolated following methods previously described (43). In brief, cells were washed 2x with ice-cold PBS and then incubated for 10 min on ice with swelling buffer (10mM Tris-HCl [pH 7.5], 10% glycerol, 3mM CaCl_2_, 2mM, MgCl2_2_, 0.5mM DTT, protease inhibitors [Roche] and 4 U/ml RNase inhibitors [RNaseOUT, ThermoFisher]). Cells were scraped from the plate, pelleted via centrifugation at 600 x g for 10 min (4°C), resuspended in lysis buffer (swelling buffer + 0.5% Igepal) and then incubated on ice for 20 min to release nuclei. Nuclei were pelleted by centrifugation at 1500 x g for 5 min (4°C), washed 2x with lysis buffer followed by a final wash in storage buffer (50mM Tris-HCl [pH 8.0], 25% glycerol, 5mM MgAcetate, 0.1mM EDTA, 5mM DTT). Nuclei were resuspended in storage buffer, flash frozen in LN_2_ and stored at -80°C.

### Precision nuclear run-on assay

The nuclear run-on assays were in performed in triplicate as described in (21,43,44) for Mutu-I and Akata experiments and in duplicate for the LCL experiment. Frozen nuclei were thawed on ice and S2 nuclei spiked-in to Mutu-I/Akata/LCL cells at a ratio of 1:1000. An equal volume of run-on buffer (10mM Tris-HCl [pH 8.0], 5mM MgCl_2_, 1mM DTT, 300 mM KCl, 20µM biotin-11-ATP, -UTP, -GTP, -CTP, 0.4 U/ml RNase inhibitor [RNaseOUT ThermoFisher] and 1% Sarkosyl) was added to thawed nuclei. Nuclear run-on was performed under constant shaking for 3 min on a vortex shaker at 37°C. The reaction was terminated by the addition of TRIzol LS (ThermoFisher) and vortexed to homogenize.

### Library preparation

RNA was extracted from run-on nuclei via TRIZOL extraction. Limited hydrolysis of RNA was done by the addition of 0.2N NaOH and incubation on ice for 20 min. Unincorporated nucleotides were removed using a P-30 column (Bio-Rad). Biotinylated RNA was purified using streptavidin M280 Dynabeads (ThermoFisher) after a series of washes. Bead-bound RNA was washed 2x in ice-cold high salt wash (50mM Tris-HCl [pH 7.4], 2M NaCl, 0.5% Triton X-100, 0.4 U/ml RNase inhibitor [RNaseOUT ThermoFisher]), 2x in ice-cold medium salt wash (10mM Tris-HCl [pH 7.4], 300mM NaCl, 0.1% Triton X-100, 0.4 U/ml RNase inhibitor [RNaseOUT ThermoFisher]) and once in ice-cold low salt wash (5mM Tris-HCl [pH 7.4], 0.1% Triton X-100, 0.4 U/ml RNase inhibitor [RNaseOUT ThermoFisher]). RNA was eluted from beads with two TRIzol extractions.

The 3’-RNA adapter with a 5’ phosphate (5’-Phos) and 3’ inverted dT (InvdT) (5’-Phos-GAUCGUCGGACUGUAGAACUCUGAAC-3’-InvdT) was ligated to the 3’ end of the RNA using T4 RNA ligase I (NEB). RNA was purified as above by binding to streptavidin beads, washing and TRIzol extracted followed by removal of the 5’ cap with 10 U of 5’-pyrophosphohydrolase (RppH) (NEB) and 5’ end repair with T4 PNK (NEB). The 5’-RNA adapter (5’-CCUUGGCACCCGAGAAUUCCA-3’) was ligated to the 5’ end of the RNA using T4 RNA ligase I (NEB). RNA was purified as above by binding to streptavidin beads, washing and TRIzol extraction.

RNA was reverse transcribed with SuperScript III reverse transcriptase (ThermoFisher) using the RNA PCR primer 1 (5’-AATGATACGGCGACCACCGAGATCTACACGTTCAGAGTTCTACAGTCCGA-3’). The cDNA was PCR amplified with Phusion high-fidelity DNA Polymerase (NEB) using barcoded Illumina PCR index primers. Libraries were purified on an 8% polyacrylamide-TBE gel and assessed on a fragment analyzer (Bioanalyzer) to confirm correct library size. Note: one library replicate of Mutu-I 1h reactivated did not meet quality requirements and was not analyzed. Libraries were sequenced on an Illumina NextSeq 500, performed by the GeneLab at Louisiana State University School of Veterinary Medicine, or by the Biotechnology Resource center (BRC) Genomics facility *(RRID:SCR_021727)* at the Cornell Institute of Biotechnology.

### Read processing

FastQ files were processed using the PRO-Seq pipeline developed by Danko and colleagues at Cornell University https://github.com/Danko-Lab/utils/tree/master/proseq. The pipeline first trims adapters and aligns reads to a concatenated genome file containing hg38, dm3, EBV genomes and rRNA. The EBV genome used was dependent on the experiment. (Mutu-1: KC207814.1, Akata: KC207813.1 and LCL: NC_007605.1.) SeqMonk software (45) was used to align the output .bam files to the drosophila, EBV and Hg38 genomes. For viral genes that overlap on the same strand, assignment of the originating gene was not possible. Read counts in these instances were therefore assigned to all overlapping genes. Drosophila spike-in normalization was used to account for variation in sequencing depth between libraries. Libraries were normalized relative to the library with the largest drosophila read count, with the scale factor set to 1. The total drosophila reads for all libraries were divided from the largest number to determine the relative normalization scale factor. The IGV genome browser (23) was used for visualization of the output .bw files (containing only the 3’ final read position). Individual reads for each nucleotide across the EBV genome was extracted from the .bw files using multiBigWigSummary from deepTools (46) and normalized as described above. The resulting .txt files were used for analysis in Seqmonk and converted to .bedgraph for IGV visualisation for figure production.

### Fold change analysis

Fold change analysis was performed using the R package DeSeq2 (25). DeSeq2 for EBV genes was performed incorporating Hg38 genes to account for the library size correction step. Full viral gene DeSeq2 fold change and p-values are given in Appendix 2.

### dREG analysis

The PRO-Seq output .bw files were merged for each replicate using the mergeBigWigs script from the Danko PRO-Seq pipeline https://github.com/Danko-Lab/utils/tree/master/proseq. The merged .bw files were then imported into dREG web-based server, freely available at https://dreg.dnasequence.org/ and dREG analysis performed (24). To identify overlapping regions between cell lines, peaks were aligned to a consensus reference genome (GenBank NC_007605.1). Full list of dREG significant peaks and scores are given in Appendix 1.

### ChIP-Seq and ATAC-Seq data

WIG files of ChIP-Seq data from the EBV genome were downloaded from the EBV portal (7), freely available at https://ebv.wistar.upenn.edu/. ATAC-Seq data is available from the NCBI GEO database under accession GSE172476.

### Chromatin immunoprecipitation (ChIP)-qPCR Assays

ChIP-qPCR assays were performed as described previously (47). Quantification of precipitated DNA was determined using real-time PCR and the delta Ct method for relative quantitation (ABI 7900HT Fast Real-Time PCR System). Rabbit IgG (Cell Signaling, 2729S), rabbit anti-CTCF (EMD Millipore, 07-729) and rabbit anti-Rad21 (Abcam, ab992) were used in ChIP assays. Primers for ChIP assays are listed in Table S3

### RT-qCPR

The EBV WT and ΔCTCF LCLs treated with DMSO or lytic activating compound C60 (28) were harvested at indicated time points. Total RNA was isolated using the RNeasy kit (Qiagen) and then treated with DNase I by using the DNase treatment and removal kit from Ambion. Reverse transcription was carried out on equal amounts of DNase-treated RNA using SuperScriptIV reverse transcriptase (Invitrogen), random priming mix (New England Biolabs), and RNase inhibitor (New England Biolabs) following the manufacturer’s instructions. qPCR was performed with Power SYBR green 2x PCR mastermix with primers (Table S3). Data was normalized to the gene GUSB.

## Data availability

The PRO-Seq data will be publicly available upon publication on the GEO database under the accession numbers: GSE188584 and GSE211247. Reviewers can access the data using secure access token itmzsuokdjqnvif and argvgsqcnjmprov, respectively.

## Acknowledgements

Portions of this research were conducted with the high-performance SuperMic supercomputer provided by Louisiana State University (http://www.hpc.lsu.edu) and we thank Dr Le Yan for his help in maintaining the PRO-Seq pipeline.

We thank Thaya Stoufflet and Dr. Vladimir Chouljenko at the LSU SVM GeneLab for their help with the NextSeq 500 sequencing. We also thank Dr. Claire Birkenheuer for all her assistance and discussion with the PRO-Seq technique.

This work was supported by NIH grants R01 AI 141968 and R21 AI 148926 to JDB and R01 DE017336 and R01 CA093606 to PML

## Supporting Information

**Figure S1: RT-qPCR validation of EBV reactivation.**

Relative expression of EBV transcripts in untreated and reactivated cells. (A) Mutu-I cells untreated and after 24 h treatment with NaB/TPA. (B) Akata cells untreated and after 4 and 24 h treatment with anti-IgG.

**Figure S2: PRO-Seq peaks do not align consistently with known canonical or novel transcription start sites.**

High resolution IGV view of PRO-Seq tracks from EBV genome regions with the most significant increase in Pol activity during early reactivation in Mutu-I cells. (A) EBER region. (B) EBER region, scaled to improve visualization. (C) BCRF1-Cp region. (D) LMP-2A, exon 1. (E) RPMS1 region. (F) The BZLF1/BRLF1, shown for comparison. Data shown is the mean of 2-3 biological replicates, scaled relative to a drosophila spike-in. EBV GenBank annotation taken from accession number: KC207814.1 and EBV novel transcripts taken from O’Grady et al. (25). Solid red/orange indicates coding regions. Black lines indicate non-coding regions of transcripts. Transcription start sites regions are highlighted in green.

**Figure S3: Distribution of active polymerase on the EBV episome in latent and reactivated Akata cells.**

(A) IGV view of the distribution of PRO-Seq reads along the EBV episome in latent Akata cells and at 1, 4 and 12h post-reactivation with anti-IgG (mean of 3 biological replicates, scaled relative to drosophila spike-in). (B) IGV view of the distribution of PRO-Seq reads 24h post-treatment with DMSO (latency) or C60 (reactivation) along the EBV episome of a WT LCL and the ΔCTCF LCL EBV mutant. The EBV bacmids used to generate these LCLs were generated in B95-8, which contains a deletion within the shaded area. Data presented is the mean of 2 biological replicates, scaled relative to drosophila spike-in. Fold increase (above 0.5) of normalised PRO-Seq reads (in 50bp bins) in C60 relative to DMSO treatment across the EBV episome is shown in green

**Table S1: Total normalized read counts from Mutu-I Pro-Seq experiment**

**Table S2: Total normalized read counts from Pro-Seq experiment**

**Table S3: Total normalized read counts from LCL Pro-Seq experiment.**

**Table S3: PCR primers used for PCR**

**Appendix 1: Details of significant dREG peaks in Mutu-I and Akata cells**.

**Appendix 2: EBV genes DeSeq2 results**.

