## Supplementary figures and images for "Reactivation of Epstein Barr Virus from Latency Involves Increased RNA Polymerase Activity at CTCF Binding Sites on The Viral Genome"

### Supplemental Figure 1

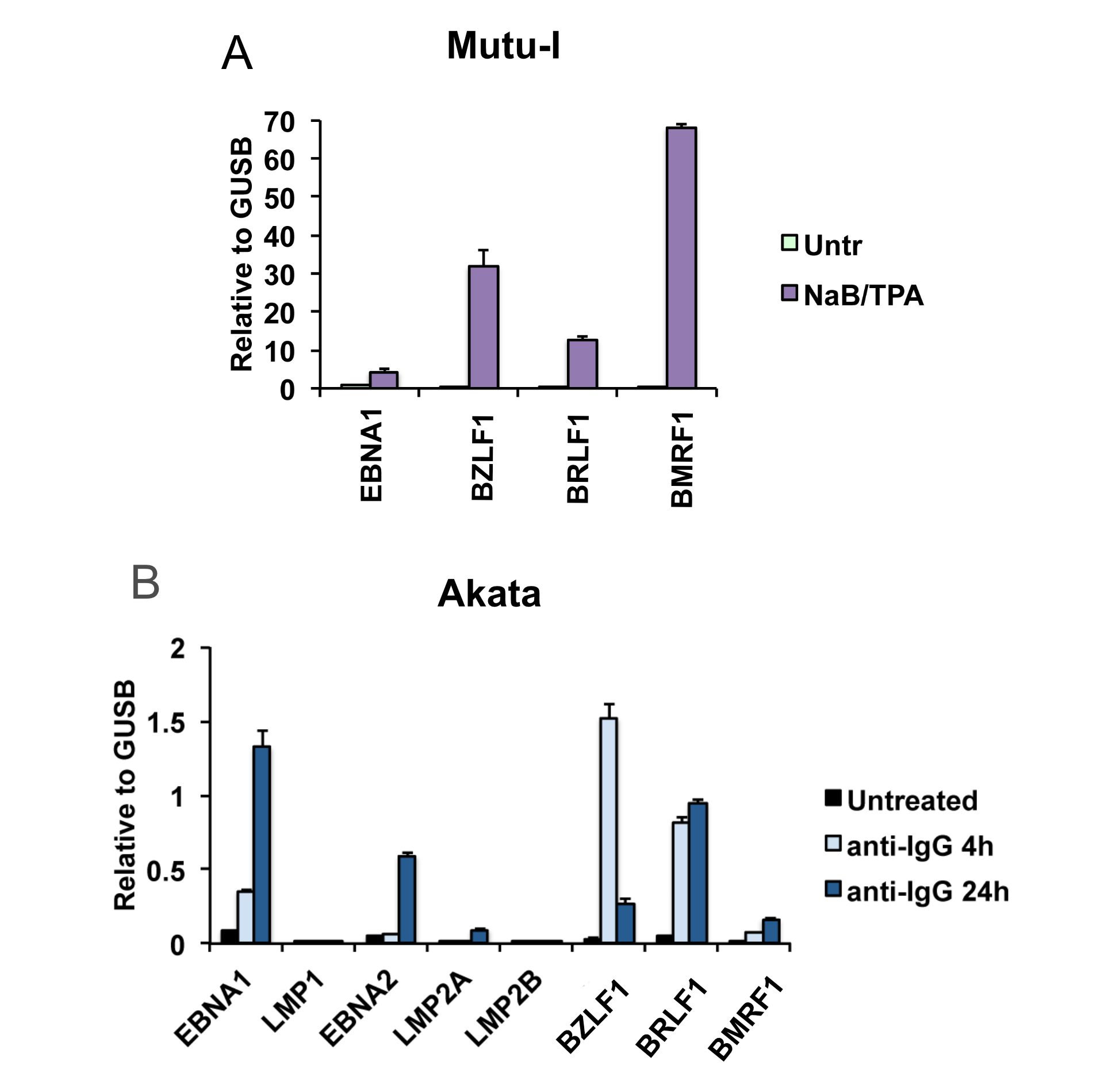

### Supplemental Figure 2

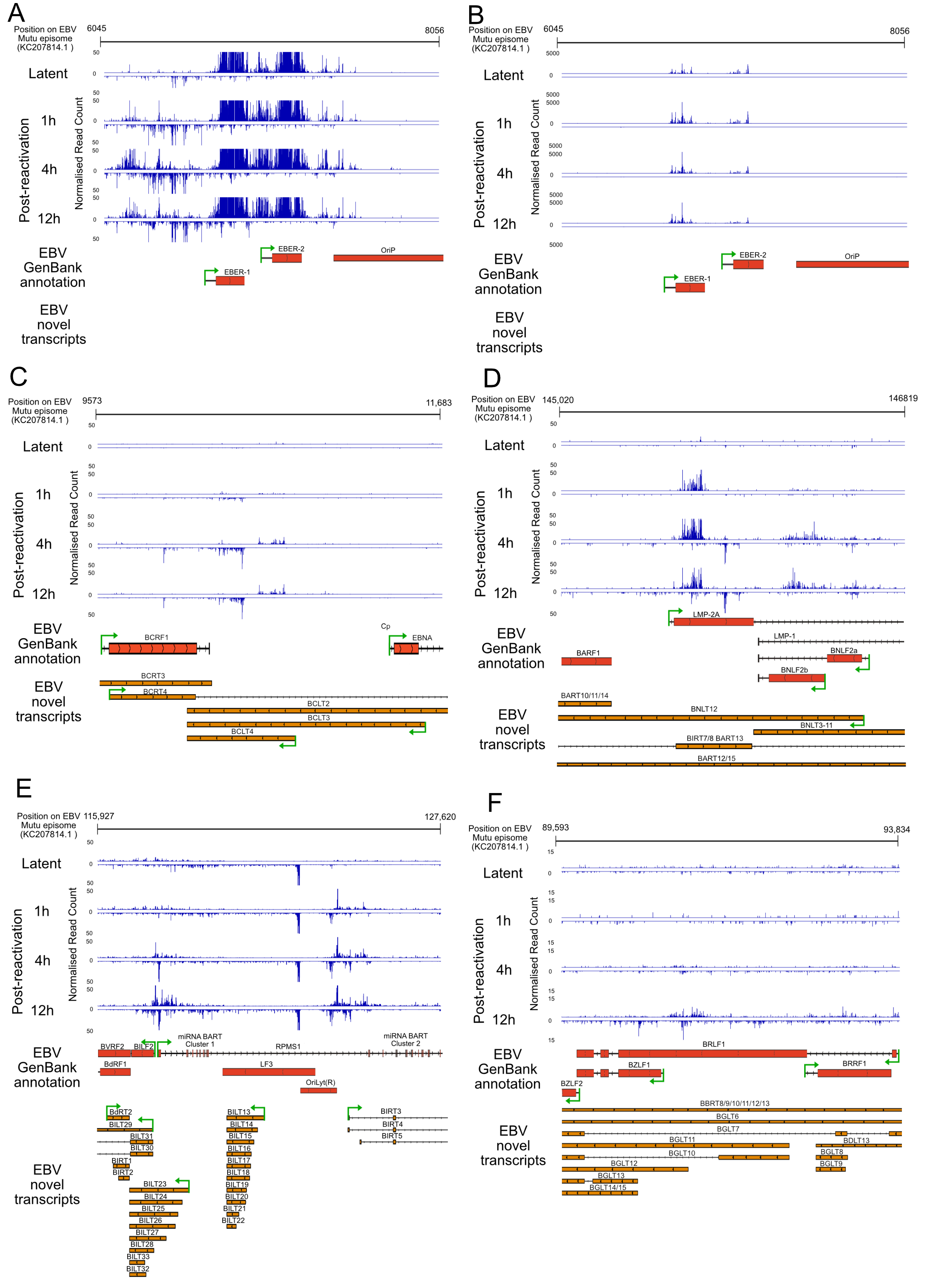

### Supplemental Figure 3

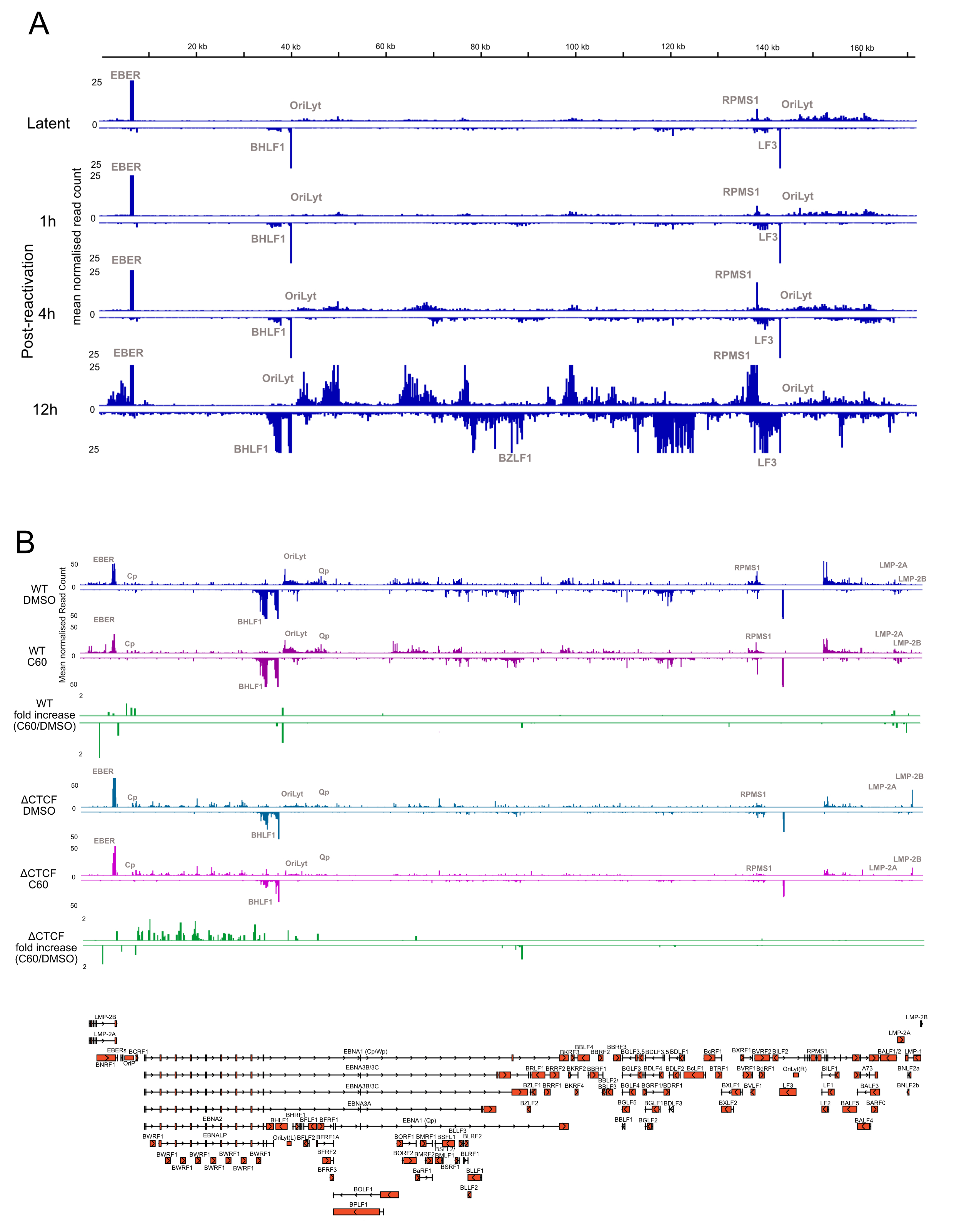
