## Supplemental Tables 1-3 for "Reactivation of Epstein Barr Virus from Latency Involves Increased RNA Polymerase Activity at CTCF Binding Sites on The Viral Genome"

Table S1: Total normalized read counts from Mutu-I PRO-Seq experiment

| | **Total normalized reads aligning to EBV genome** | | | | **Total normalized reads aligning to Hg38 genome** | | | |
|---|---|---|---|---|---|---|---|---|
| | **replicate** | | | | **replicate** | | | |
| **Time** | **1** | **2** | **3** | **mean** | **1** | **2** | **3** | **mean** |
| Latent | 70701.30 | 91995.10 | 86629.70 | 85479.0 (±12054.6) | 53831097.67 | 53097782.20 | 54554179.63 | 50215601.10 (±728204.70) |
| 1h post-reactivation | 109713.90 | 138352.79 | Did not pass QC | 127096.9 (±20709.2) | 56350362.75 | 56807777.67 | Did not pass QC | 56579070.21 (± 323441.19) |
| 4h post-reactivation | 102712.70 | 122201.85 | 115782.23 | 118101 (±10026) | 54042412.30 | 58437175.11 | 56367206.50 | 56282264.64 (±2198612.37) |
| 12h post-reactivation | 143867.00 | 143304.49 | 135207.47 | 146689.3 (±4573.7) | 55283366.00 | 54743156.95 | 50215601.10 | 53414041.35 (± 2783068.717) |

Data is normalized to drosophila spike-in. ± is standard deviation of the mean.

Table S2: Total normalized read counts from Akata PRO-Seq experiment

| | **Total normalized reads aligning to EBV genome** | | | | **Total normalized reads aligning to Hg38 genome** | | | |
|---|---|---|---|---|---|---|---|---|
| | **replicate** | | | | **replicate** | | | |
| **Time** | **1** | **2** | **3** | **mean** | **1** | **2** | **3** | **mean** |
| Latent | 9359.04 | 7511.51 | 10384.05 | 9026.2 (±1453.0) | 65935314.82 | 66743793.44 | 65915329.10 | 66457058.6 (±398303.3) |
| 1h post-reactivation | 7778.70 | 7451.82 | 9940.57 | 8405.6 (±1377.0) | 62660324.02 | 61762504.45 | 61351575.29 | 62929900.7 (±568982.3) |
| 4h post-reactivation | 14359.93 | 14067.00 | 17133.56 | 15075.3 (±1671.5) | 63701916.25 | 63756813.00 | 63791986.94 | 63783354 (±29349.7) |
| 12h post-reactivation | 69665.94 | 73802.61 | 78467.97 | 73258.5 (±4432.3) | 63820794.41 | 64408264.12 | 62143552.58 | 63395247.2 (±1069515.9) |

Data is normalized to drosophila spike-in. ± is standard deviation of the mean.

Table S3: Total normalized read counts from LCL Pro-Seq experiment

| | **Total reads aligning to EBV genome** | | | | | | **Total reads aligning to Hg38 genome** | | | | | |
|---|---|---|---|---|---|---|---|---|---|---|---|---|
| **Treatment** | **WT LCL** | | | **ΔCTCF LCL** | | | **WT LCL** | | | **ΔCTCF LCL** | | |
| | **1** | **2** | **mean** | **1** | **2** | **mean** | **1** | **2** | **mean** | **1** | **2** | **mean** |
| DMSO | 570421.75 | 429688.09 | 500054.9 (±99513.7) | 194696.15 | 164828.00 | 179762.1 (±21119.9) | 118238633 | 85874578.8 | 102056606 (±22884842) | 74704585 | 60510364 | 67607474.5 (±10036829.9) |
| C60 | 440841.33 | 426857.75 | 433849.5 (±9887.9) | 190282.02 | 187306.40 | 188794.2 (±2104.1) | 106626454 | 100222831 | 103424642 (±4528045.75) | 61921071.7 | 67879071.1 | 64900071.4 (±4212941.77) |

Data is normalized to drosophila spike-in. ± is standard deviation of the mean.
