## Supplemental Table 4 for "Reactivation of Epstein Barr Virus from Latency Involves Increased RNA Polymerase Activity at CTCF Binding Sites on The Viral Genome"

Table S4: Primers used in this study

| Primer | Sequence | Use |
|---|---|---|
| LMP_CTCF_BS_5' | TATACGAAGAAGCGGGCAGAGGAA | ChIP-qCPR |
| LMP_CTCF_BS_3' | TGACCTGTTGTCCCTGAGATGTGA | ChIP-qCPR |
| Cp_CTCF_BS_5' | ACATTTTGTTCTAGGTCCATCTTAGGA | ChIP-qCPR |
| Cp_CTCF_BS_3' | TGCCCTCCCCACTTCTCTT | ChIP-qCPR |
| Qp_CTCF_BS_5' | AAATTGGGTGACCACTGAGGGAGT | ChIP-qCPR |
| Qp_CTCF_BS_3' | ATAGCATGTATTACCCGCCATCCG | ChIP-qCPR |
| LMP1p_5' | ACGTCAGAGTAACGCGTGTTTC | ChIP-qCPR |
| LMP1p_3' | GCAGACCCCGCAAATCC | ChIP-qCPR |
| XqYq_CTCF_5' | GGTGGAACTTCAGTAATCCGAAA | ChIP-qCPR |
| XqYq_CTCF_3' | AGCAAGCGGGTCCTGTAGTG | ChIP-qCPR |
| 10q_CTCF_5' | ACGGTGCTCTGCCATTGC | ChIP-qCPR |
| 10q_CTCF_3' | GGCGCTGGACACCACTGTA | ChIP-qCPR |
| GPR56_5' | GTCCTTGTCCCCAGCAGAAA | ChIP-qCPR |
| GPR56_3' | CCCAGGGTCCAGAGCATCT | ChIP-qCPR |
| GAPDH_5' | CGGTGCGTGCCCAGTT | ChIP-qCPR |
| GAPDH_3' | CTACTTTCTCCCCGCTTTTTTT | ChIP-qCPR |
| GUSB_5’ | CGCCCTGCCTATCTGTATTC | RT-qPCR |
| GUSB_3’ | TCCCCACAGGGAGTGTGTAG | RT-qPCR |
| BZLF1_5’ | TCTGAACTAGAAATAAAGCGATACAAGAA | RT-qPCR |
| BZLF1_3’ | TTGGGCACATCTGCTTCAAC | RT-qPCR |
| BMRF1_5’ | TTGGGCAGGTGCTGTTGAT | RT-qPCR |
| BMRF1_3’ | TGCCCACTTCTGCAACGA | RT-qPCR |
| LMP2A_5’ | TTCAGGCCGGTTTTGCA | RT-qPCR |
| LMP2A_3’ | GCCCATTGGCACCATTTCTA | RT-qPCR |
| EBER1_5’ | TTTGCTAGGGAGGAGACGTGTGT | RT-qPCR |
| EBER1_3’ | AAGCAGAGTCTGGGAAGACAACCA | RT-qPCR |
| EBER2_5’ | TTGCCCTAGTGGTTTCGGACACA | RT-qPCR |
| EBER2_3’ | ACTTGCAAATGCTCTAGGCGGGAA | RT-qPCR |
